# Analyzing single-molecule dynamics with both complex types of motion and complex transition kinetics: Benchmarking of ExaTrack

**DOI:** 10.64898/2026.01.22.700663

**Authors:** François Simon, Chris H. Wiggins, Lucien E. Weiss

**Affiliations:** Department of Applied Physics and Applied Mathematics, Columbia University, New York City, USA; Department of Engineering Physics, Polytechnique Montréal, Montréal, Québec, Canada

**Keywords:** Single-particle tracking, Multi-state model, Diffusive motion, Directed motion, Confined motion, non-Markovian transition kinetics

## Abstract

Single-particle tracking (SPT) is a tool of growing importance which enables biologists to better understand the dynamics of protein interactions at the single-molecule level and *in vivo*. However, the stochastic nature of the motion of single molecules, the wide variety of types of motion that they can experience and complex transition kinetics between the different states of motion are challenging factors that complicate the interpretation of SPT data. This article presents and benchmarks the tool ExaTrack. Like previous tools, it can handle particles moving in one or multiple states of motion with transitions between states. Its unique feature is that it can simultaneously handle a wide range of complex types of motion such as diffusive motion, directed motion and confined motion while also managing a variety of transition kinetics such as memoryless first-order transitions or more complex time-dependent state transitions. This manuscript focuses on the benchmarking of ExaTrack on simulated data.

## Introduction

Single-particle tracking (SPT) is a powerful approach for studying the behaviors of single molecules and reveal microenvironments in cells. Recent technological advances in camera sensitivity, fluorescent dye brightness and stability, and specific labeling strategies have enabled single-molecule assays to become increasingly routine. An important aspect of SPT pipelines is the analysis step to make sense of the ensemble track behaviors. This is challenging due to the stochastic nature of motion on the mesoscale and local inhomogeneity in cells. Measured heterogeneous behaviors can therefore be caused by a multitude of factors including spatially varying viscosity, obstacles (1–5), elasticity, and molecular-interaction partners. These interactions between tracked particles and targets can cause transitions between types of motion, for instance, between free diffusion and a bound state. Such behaviors reveal interactions with a larger substrate (6–9). Many of these reactions have been found to follow first-order kinetics, and a range of strategies have been developed to recover the parameters governing the states of motion and Markovian (memoryless) transition rates between them (10–14). However, the aforementioned Hidden Markov models are constrained to Brownian diffusion.

Oftentimes the particle motion encountered in cells is more complex than pure Brownian diffusion. For instance, particles, such as molecular motors, can move in a directed manner (15, 16). The diffusion of some particles can also be confined to subcellular compartments. Non-Brownian motion is often referred to as anomalous motion (17, 18) and can arise from a variety of motion models. To compare and determine the most likely motion model, machine-learning-based tools, trained on simulated data, have become an increasingly popular approach (19–22). However, most machine learning techniques deployed for SPT, so far, are based on machine-learning architectures that require a large number of parameters (CNN, RNN, transformers, etc.) and therefore risk overfitting and low generalizability (23, 24). While some methods consider hidden variable (unobserved properties of the molecule) to model non-Brownian diffusion and state transitions, they are available for either confined motion (25, 26) or directed motion (27).

Here, we present a probabilistic tool to extract parameters from particles in multiple states of motion that can be diffusive, directed or confined with state transitions within tracks. In our method, called ExaTrack, each state of motion is modeled by two hidden variables, a particle’s position and an anomalous-diffusion variable which represents either a velocity vector in directed motion or a potential well for confined motion (Fig. 1a). To model transitions, ExaTrack considers transitions between the different states of motion and these transitions with gamma-distributed lifetimes. This framework allows transitions to follow either a memoryless Markov transition model or a more time-dependent model. ExaTrack can be used 1) to determine the number of states, 2) to quantify the parameters associated with each state and the transition kinetics, and 3) to label each track with their corresponding probabilities of being in each state (Fig. 1b). This preprint explains the principle of ExaTrack and benchmarks its performance by evaluating model parameter estimation and state prediction on simulated tracks across a wide range of parameters.

**Fig. 1.**
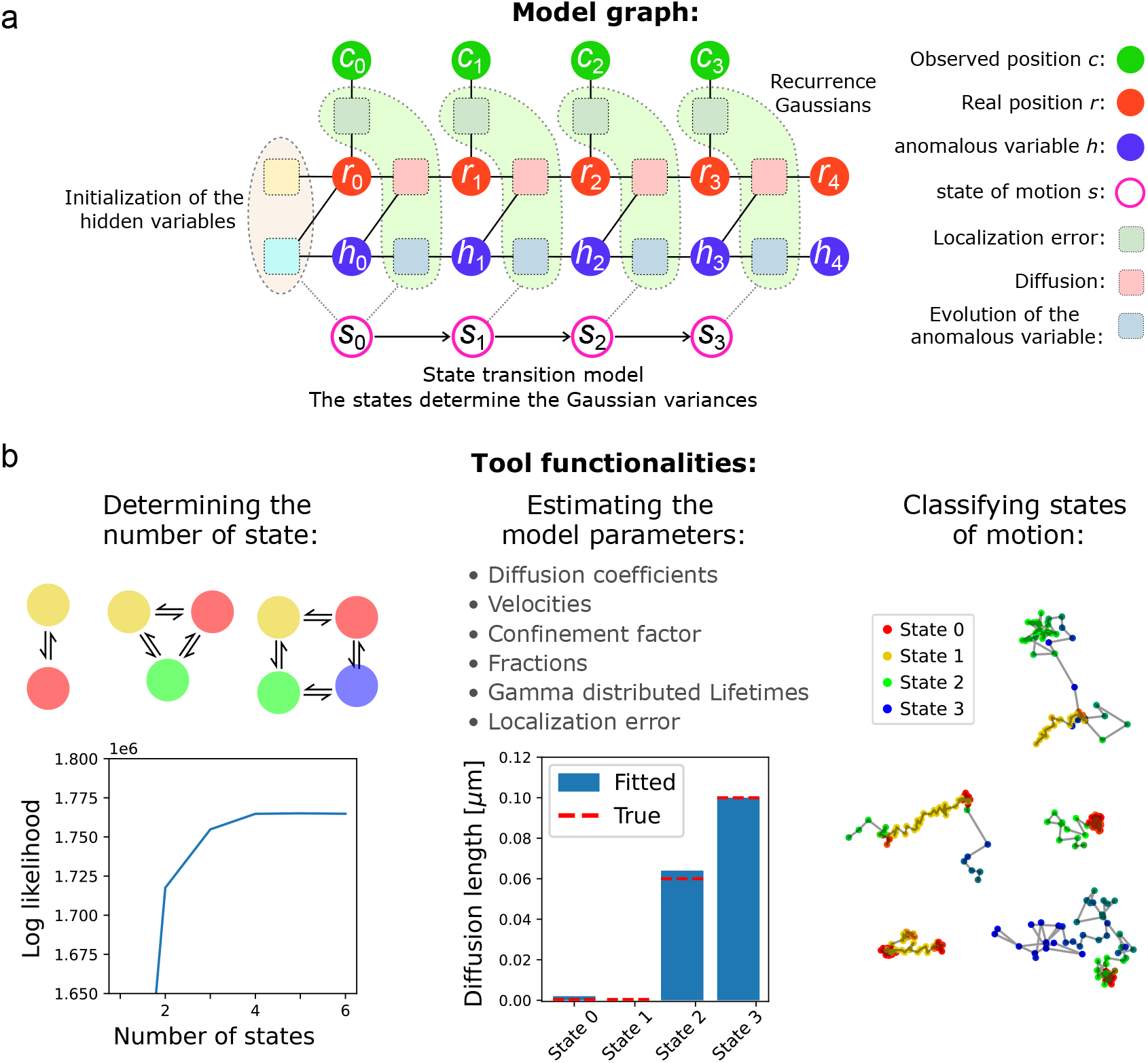
Model features. **a**: Graph representing the ExaTrack model. The probability of the track is computed from the joint probability density function by integration over the possible real positions and anomalous variables that either represent the velocity vector or the potential well and by sum over the possible sequences of states. **b**: Functionalities of ExaTrack: Determination of the number of states from the likelihood plateauing or Bayesian Information Criterion (BIC). Estimation of the model parameters. State classification of tracks.

## Results

### A. Model overview

The motion model of ExaTrack is based on our previous algorithm aTrack (24). Each state of motion is modeled by discrete updates of the particle’s true position with a diffusion sub-step and an anomalous sub-step, where the position evolves according to a random variable that either represents the velocity of the directed motion for directed motion states or the center of a potential well for confined motion states (Fig. 1a). The velocity and the potential well position are also allowed to change through time. Then localization error is taken into account by considering the relationship between the true particle position and the measured track position. For a segment of a given state of motion, the joint-probability of the track 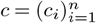, the real particle positions 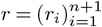, and either the velocity vector or the potential well center 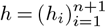 is defined as:

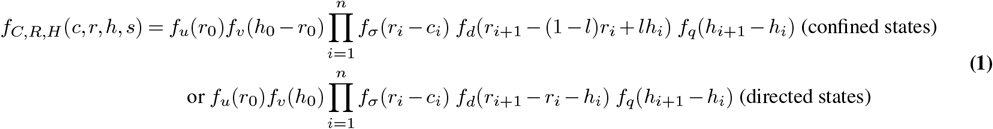

with *l* the confinement factor, *f*_*x*_ univariate Gaussians of mean 0 and standard deviation *x*, and *σ, d, q* the standard deviations of the localization error, the diffusion length and the change of anomalous variable respectively. Note that simple diffusion is included in both directed motion and confined diffusion models as a limit case when the anomalous variable has no impact. Particles can also transition from one motion state to another at any time step, either following a Markov model with exponentially distributed lifetimes, or with more complex gamma-distributed lifetimes (see Fig. 7a). To estimate the model parameters, we use the maximum likelihood estimate principle. To do so, we calculate the probability of the track by recurrent integrations over the possible continuous variables *r* and *h* and by summing over the possible sequences of states 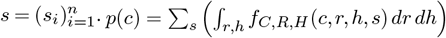. The recurrent integration is performed automatically using the same principle as (28). Considering all the possible sequences of states is challenging as the number of sequences scales with the number of states to the power of the sequence length, which makes the calculation quickly intractable as the number of states or the sequence length increases. To alleviate this issue, we found that considering a moment matching approximation that averages the coefficients and sums the probabilities of sequences that share the same previous states provides a good trade-off between speed and accuracy. See the Methods section for more details.

### B. Fitting motion and kinetic parameters for memoryless transition models

To verify that the algorithm recovers motion parameters accurately, we benchmarked ExaTrack on a range of synthetic datasets (see the Methods section for a description of track simulations) with frame times Δ*t* = 20 ms. In a first test, we simulated populations of tracks with 1 immobile state and 1 diffusive state, without directed or confined components to verify that it can correctly estimate diffusion coefficients and transition rates.

Fig 2a shows that the estimated diffusion length of the immobile state is very low with values on the order of nanometers over a wide range of transition rates and diffusion lengths of the diffusive state. Similarly, the diffusion length of the diffusive state was also accurately estimated over the range of tested conditions (Fig 2b). The localization error of the immobile state was accurately estimated too for the range of tested parameters (Fig 2d). For diffusion lengths above 0.1 µm and transition rates below 0.2 Δ*t*^−1^, the transition rates and fraction of the immobile state were also well estimated (Fig 2c,e,f).

**Fig. 2.**
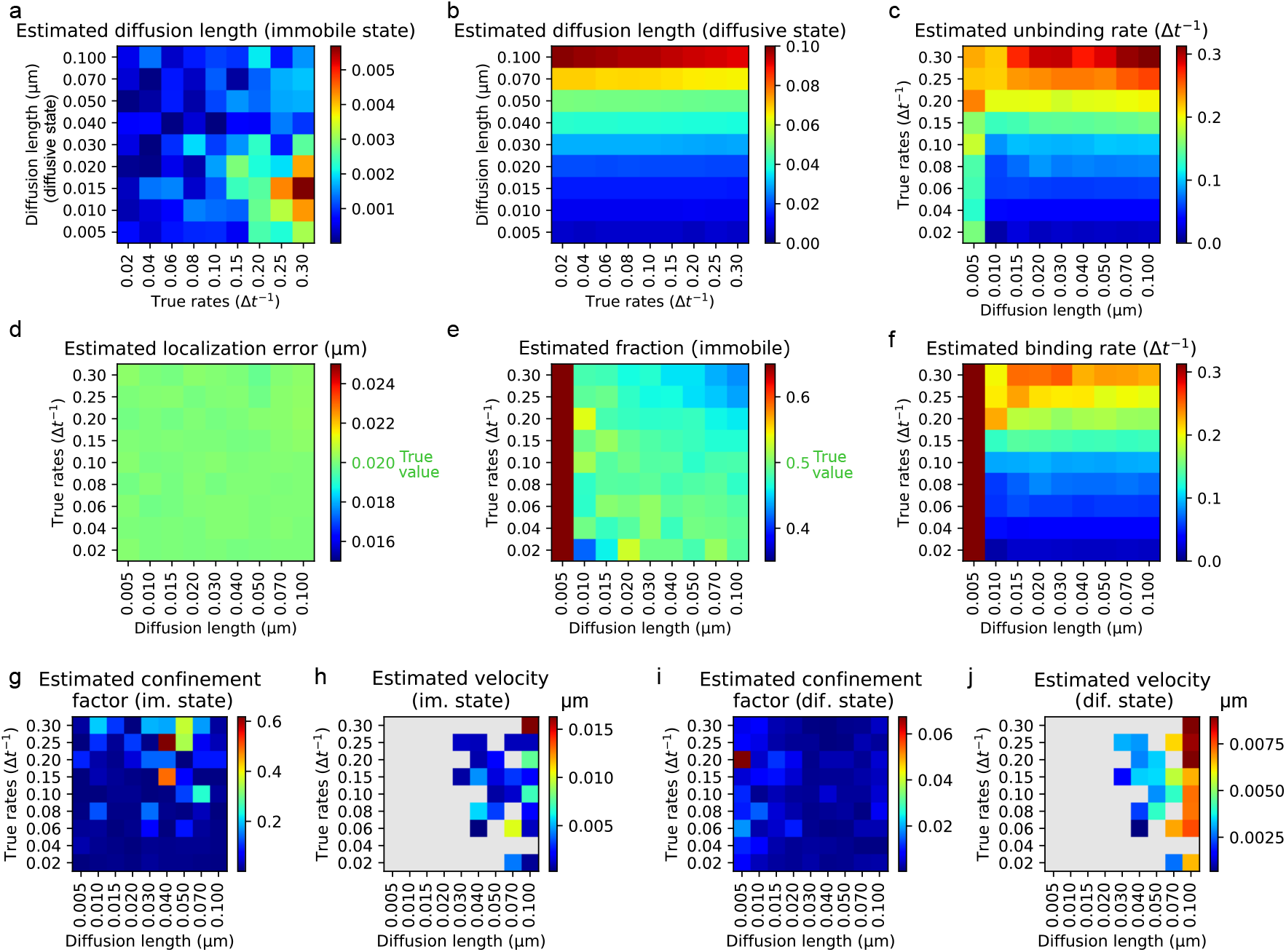
Fitting tracks with an immobile and a diffusive state. We simulated populations of 10,000 tracks with 10 time steps in two states, immobile and diffusive with continuous-time transitions between states. We then systematically varied the diffusion length of the diffusive state and the binding and unbinding transition rates (*r*_*b*_ and *r*_*u*_ respectively, with *r*_*b*_ = *r*_*u*_). The trajectories were simulated with localization errors of 0.02 µm. Each population was fitted to a 2-state ExaTrack model, 3 replicates per set of parameters. The state with the lower diffusion length was assigned as the immobile state. **a-j**: Heatmaps showing the estimated parameters of ExaTrack for combinations of diffusion lengths of the diffusive state and the transition rates (*r*_*b*_ = *r*_*u*_). **a**,**b**: The estimated diffusion lengths of the two states ( 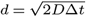 with *D* the diffusion coefficient and Δ*t* the step time). **d**: The localization error of the immobile state. **e**: The fraction of the immobile state. **c**,**f**: The unbinding and binding rates. **g-j**: Estimated confinement factor or velocity for the immobile state and for the diffusive state depending on the inferred state of motion. Grey values in the velocity heatmaps represent points where none of the 3 replicates had the state identified as directed.

As we are studying tracks without directed motion or confinement, the inferred type of motion is expected to be quite random as both confined and directed models result in very similar likelihoods. When the immobile state is identified as confined, the confinement factor can take a wide variety of values (Fig 2g). This is caused by the identifiability issue that prevents distinguishing immobile particles from particles with high confinement factor and fixed potential well. The predicted velocities of the two states and the confinement factor of the diffusive state remain low in agreement with the null underlying velocity and confinement factors(Fig 2h-j).

Next, we characterized the performance of our model for identifying directed motion and measuring velocities. To do so, we simulated tracks in either linear directed motion or diffusive motion. For all velocities higher than 0.002 µm·Δ*t*^−1^, we found that the velocity and the other parameters were reliably estimated (Fig 3). When fitting a population with an immobile state and a directed state (Fig 4), we obtained similar results, where one state is systematically estimated as directed, except that the accuracy of the estimations is lower for lower velocities.

**Fig. 3.**
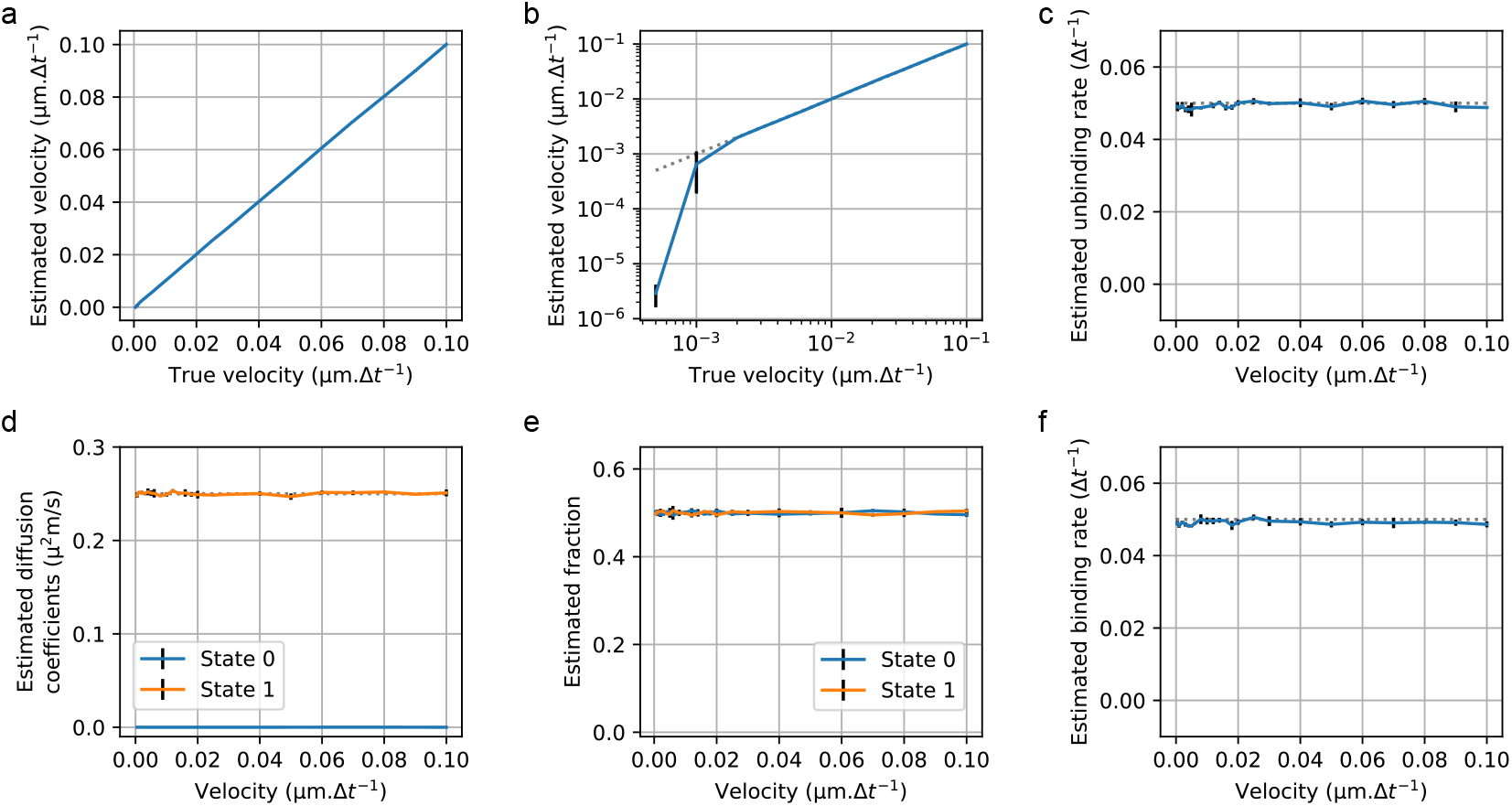
Fitting tracks with a directed (state 0) and a diffusive state (state 1). We simulated populations of 10,000 tracks of 20 time steps in two states, a directed state where particles move in linear motion of a fixed velocity and a diffusive state with continuous-time transitions between states. We then varied systematically the velocity of the directed motion to estimate its effect on our model predictions. The trajectories were simulated with localization errors of 0.02 µm. Each population was fitted to a 2-state ExaTrack model, 3 replicates per set of parameters. The curves represent the mean values of the 3 replicates and the bars represent their standard deviations. The state with the lower diffusion length was identified as the directed state. This figure presents the estimated parameters as a function of the velocity of the directed tracks. **a**,**b**: Estimated velocity as a function of the true velocity, (a) linear scale and (b) log scale to also visualize the relative changes at small velocity values. **c**,**f**: Estimated unbinding and binding rates. **d**,**e**: Estimated diffusion coefficients and fractions of the two states.

**Fig. 4.**
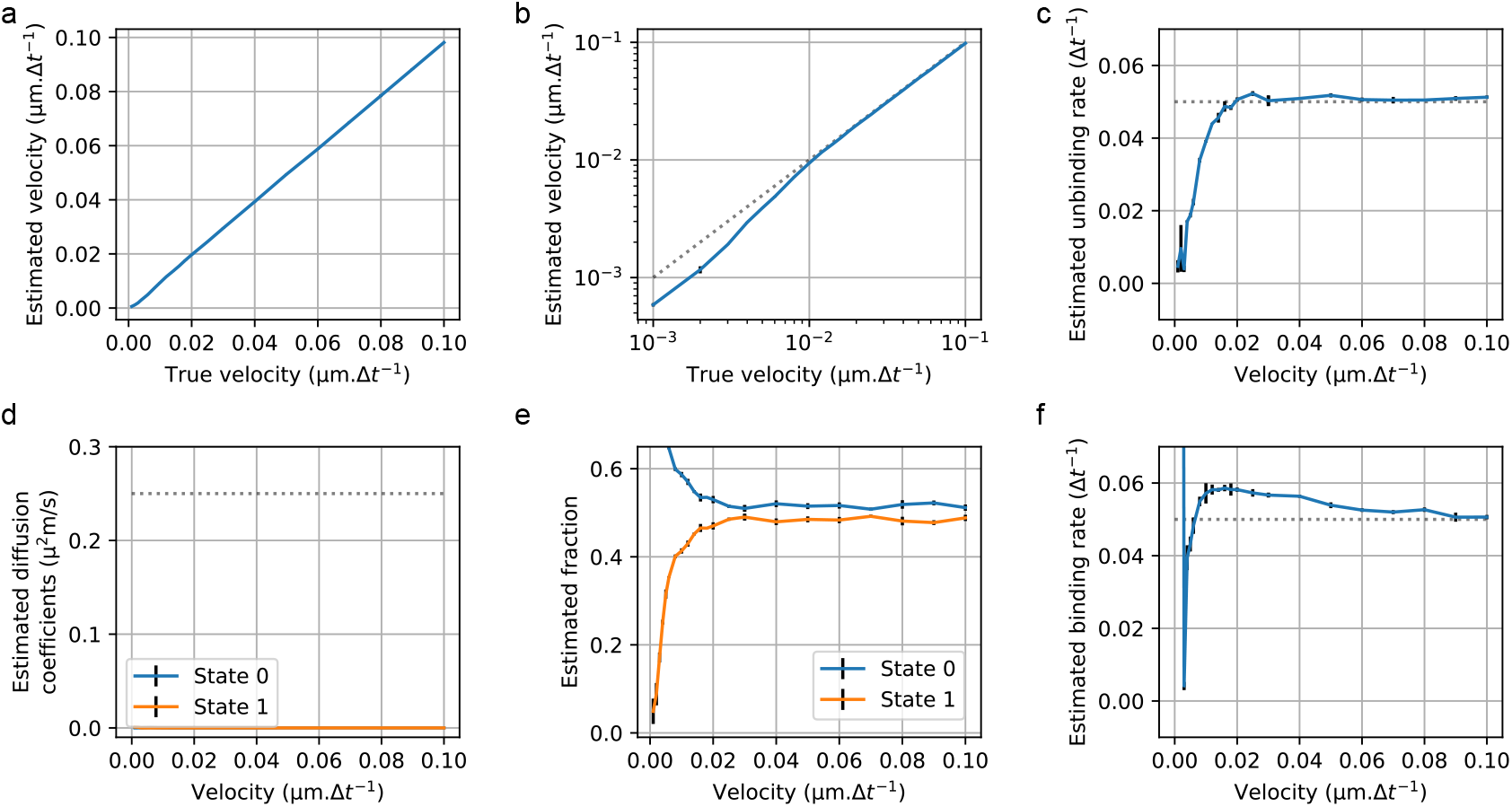
Fitting tracks with directed (state 0) and immobile states (state 1). We simulated populations of 10,000 tracks of 20 time steps in two states, a directed state where particles move in linear motion of a fixed velocity and an immobile state with continuous-time transitions between states. We then varied systematically the velocity of the directed motion to estimate its effect on our model predictions. The trajectories were simulated with localization errors of 0.02 µm. Each population was fitted to a 2-state ExaTrack model, 3 replicates per set of parameters. The curves represent the mean values of the 3 replicates and the bars represent their standard deviations. The state detected as directed with the highest velocity was identified as the directed state. This figure presents the estimated parameters as a function of the velocity of the directed tracks. **a**,**b**: Estimated velocity as a function of the true velocity, in linear scale and in log scale to also visualize the relative changes at small velocity values. **c**,**f**: Estimated unbinding and binding rates. **d**,**e**: Estimated diffusion coefficients and fractions of the two states.

To test the performance on confined tracks, we simulated a mixed population with one state either immobile (Fig 5) or diffusive (Figs 6) and a second state of confined diffusion with variable confinement factors from 0.02 to 0.99. For both experiments (Figs 5a,b and 6a,b), we found that the confined state was accurately identified as confined except for the model with a confined and a diffusive state at confinement factors of 0.99, where the confined state was identified as directed. In all other conditions, where the state was correctly identified, we found that the confinement factors were well estimated with a small positive bias of *≈*0.02 (Figs 5a). Interestingly, the transition rates were correctly identified for all the range of confinement factors, even in the case where the confined state was mistaken for a directed state (red zone in Figs 6). This is explained by the robust separation of the two states, even when the parameter estimations are incorrect. In Figs 5d and 6d, the curves of the estimated diffusion coefficients showed that the observed diffusion coefficient of the confined state decreases linearly while the confinement factor increases. This can be corrected for by dividing the diffusion coefficient by (1−*l*) as long as the confinement factor < 0.6 per. Tracks whose confined state has a confinement factor comprised between 0.6 and 0.9 results in good estimations of the confinement factor but more difficult estimations of the diffusion coefficient.

**Fig. 5.**
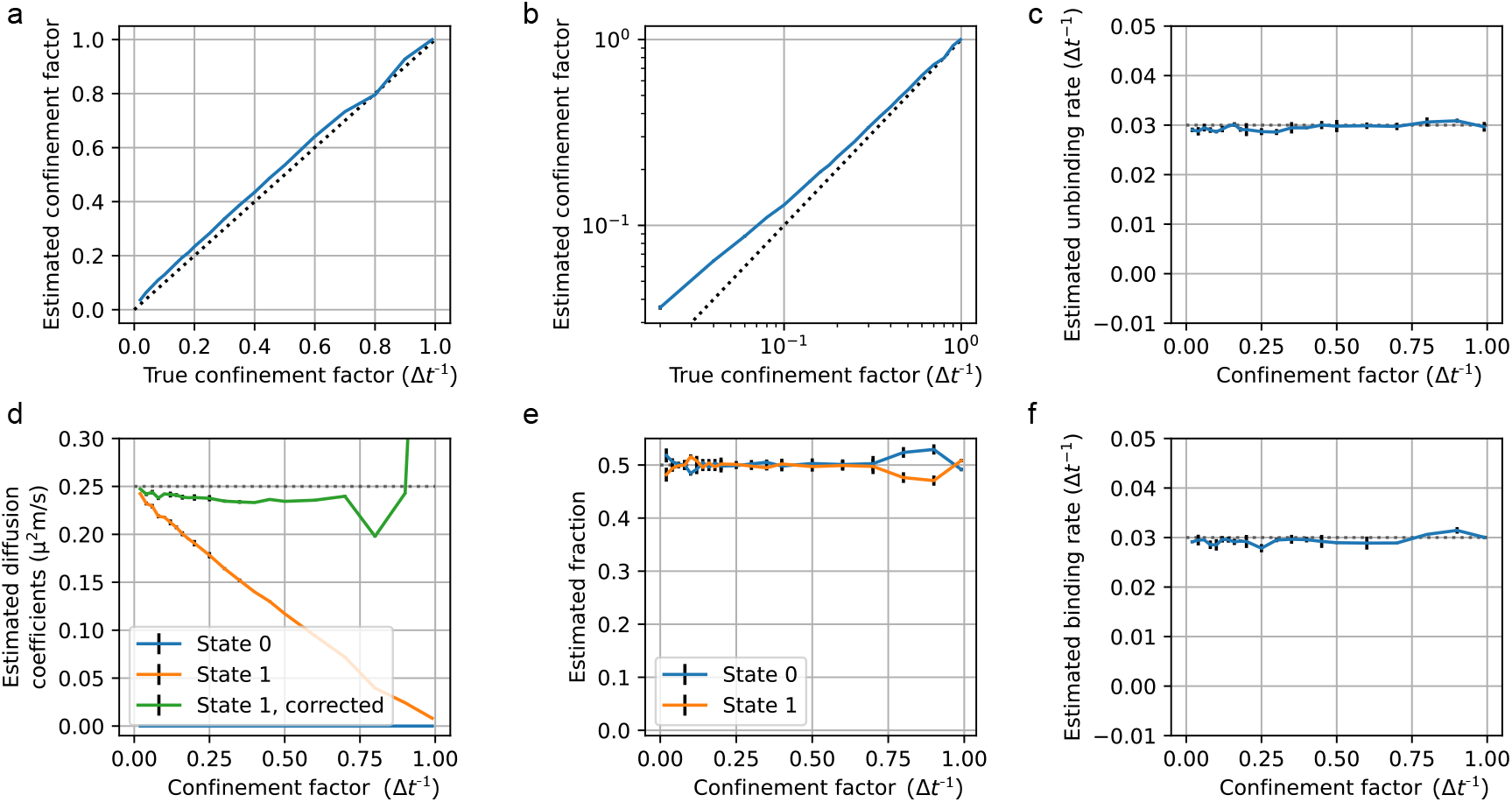
Fitting tracks with immobile (state 0) and confined states (state 1). We simulated populations of 10,000 tracks of 60 time steps in two states, a confined state where diffusive particles are stuck in a fixed potential well and an immobile state with continuous-time transitions between states. We then systematically varied the confinement factor which represents the effect of the well on the particle motion. The trajectories were simulated with localization errors of 0.02 µm. Each population was fitted to a 2-state ExaTrack model, with 3 replicates per set of parameters. The curves represent the mean values of the 3 replicates and the bars represent their standard deviations. The state with the highest confinement factor was identified as the confined state. This figure presents the estimated parameters as a function of the confinement factor of the confined state. **a**,**b**: Plots of the estimated confinement factor as a function of the true confinement factor, in linear scale and in log scale to also visualize the relative changes at small values. **c**,**f**: Plots of the estimated unbinding and binding rates. **d**,**e**: Plots of the estimated diffusion coefficients and fractions of the two states. The Curve “state 1, corrected” equals *D/*(1 *− l*) with *D* the diffusion coefficient and *l* the confinement factor.

**Fig. 6.**
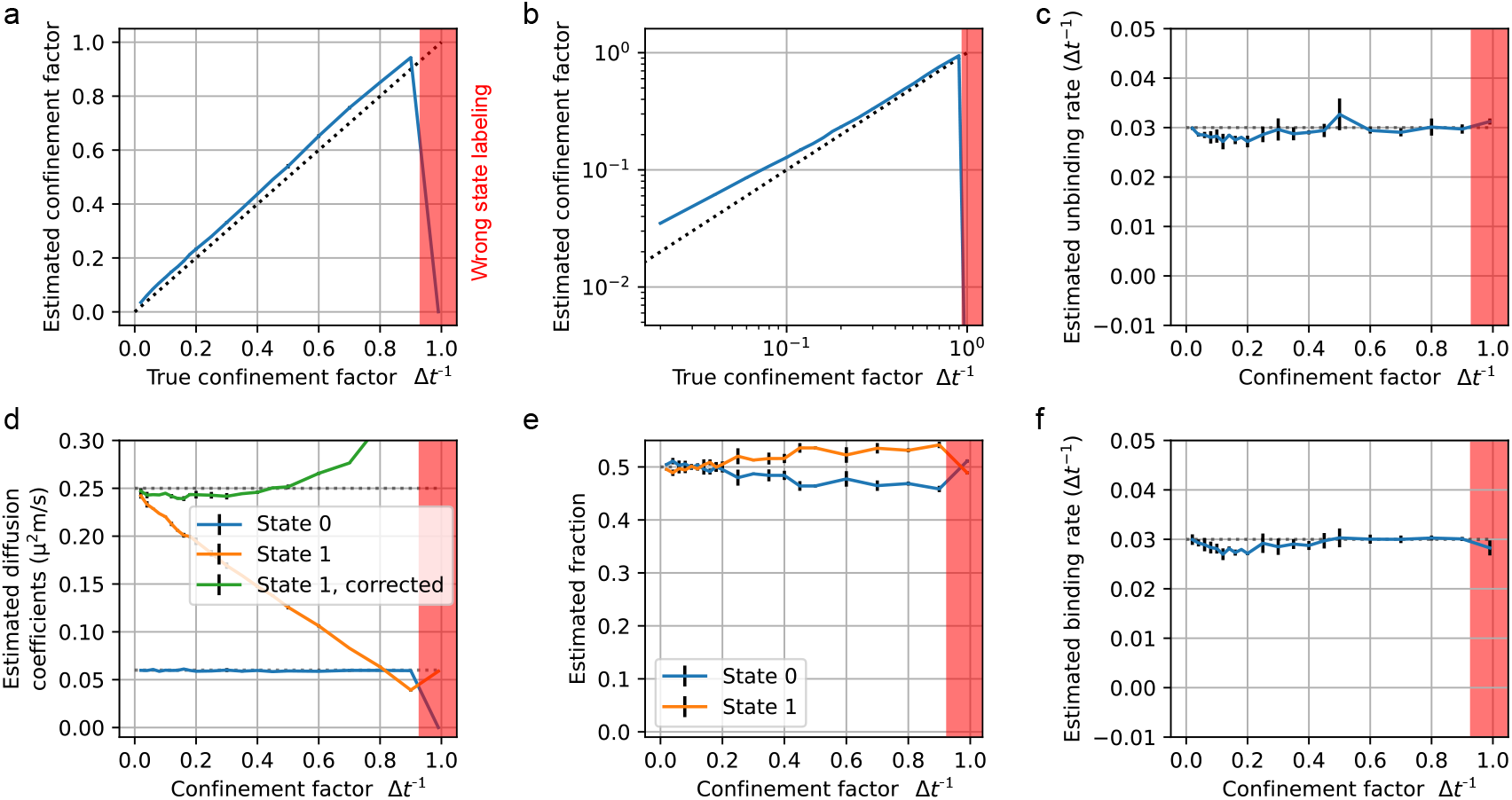
Fitting tracks with a confined (state 1) and a diffusive state (state 0). We simulated populations of 10,000 tracks of 60 time steps in two states, a confined state where diffusive particles are stuck in a fixed potential well and a diffusive state (0) with continuous-time transitions between states. We then systematically varied the confinement factor which represents the effect of the well on the particle motion. The trajectories were simulated with localization errors of 0.02 µm. Each population was fitted to a 2-state ExaTrack model, with 3 replicates per set of parameters. The curves represent the mean values of the 3 replicates and the bars represent their standard deviations. The state with the highest confinement factor was identified as the confined state. This figure presents the estimated parameters as a function of the confinement factor of the confined state. **a**,**b**: Estimated confinement factor as a function of the true confinement factor, in linear scale and in log scale to also visualize the relative changes at small values. **c**,**f**: Estimated unbinding and binding rates. **d**,**e**: Estimated diffusion coefficients and fractions of the two states.

**Fig. 7.**
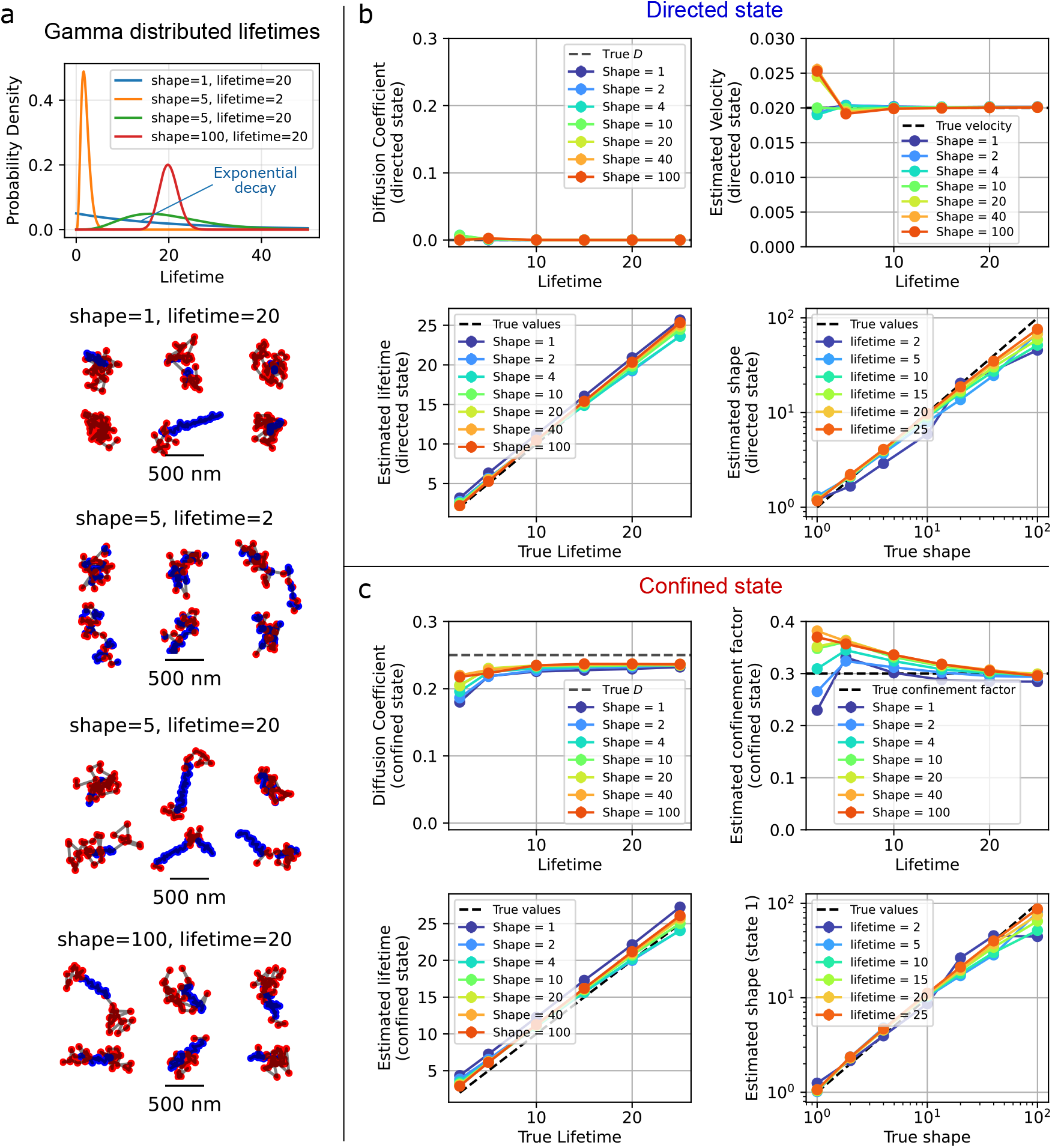
Estimating complex transition kinetics. **a**: Examples of gamma-distributed lifetimes with different shape and average lifetime parameters illustrated with tracks that follow these transition patterns where red represents the confined state, and blue the directed state. **b-c**: Results of the fittings of the parameters on 10,000 tracks of 50 time points which can be in either in a state of confined diffusion (**b**) or in a directed state of motion (**c**). The shape parameter and the average lifetime parameter were varied to verify the range where these parameters (and the others) can be estimated with high confidence. The two states of motion shared the same transition kinetics. Each data point is the mean of 3 replicates. The tracks in **a-c** are simulated with continuous transitions between the two states. The simulated confined state possesses a diffusion coefficient of 0.25 µm^2^.s^−1^ and a confinement factor of 0.3 per time step Δ*t* of 0.02 s with a fixed potential well center. The directed state has a diffusion coefficient of 0 µm^2^.s^−1^ and a velocity of 0.02 µm.Δ*t*^−1^ without changes of orientation or speed. Localization error: 0.02 µm.

### C.Modeling complex transition kinetics

To our knowledge, the currently available SPT tools that employ a transition model all assume time-independent transitions between states of motion characterized by transition rates (or probabilities) (10–12, 22, 29). There are two good reasons for this assumption: it is practical as it enables very simple Hidden Markov Models (10), and it represents the state lifetimes resulting from simple protein interactions with first-order or pseudo first-order kinetics (13, 30, 31). Nevertheless, protein-protein interactions can also be more complex with higher-order kinetics with several intermediate states (30, 32, 33) as can occur in oligomerization processes (34, 35) or for processive complexes (15, 36, 37).

Unlike first-order kinetics that result in exponentially distributed lifetimes, reactions with intermediate steps induce state life-times that are typically more localized around an average lifetime (15). A good approximation to model this type of behavior is to consider gamma-distributed lifetimes, as this distribution occurs when a reaction is composed of a sequence of sub-reactions of same kinetics (33, 38, 39) (Fig. 7)a. In order to better characterize complex transition kinetics, we integrated gamma-distributed state transitions into ExaTrack.

In Figure 7a, we show gamma-distributed lifetimes with a range of shape and lifetime parameters. A useful property of Gamma distributions is that a shape parameter of 1 corresponds to an exponential decay, the classical memoryless kinetics used in the earlier sections of this article. Higher shape parameter values, which model processes with multiple sub-steps, shift the lifetime distributions to center around the mean of the distribution and become more Gaussian-like. We tested a range of shape and lifetime parameters for simulated particle tracks to model transitions between two states of motion: a confined state and a directed state, and used ExaTrack to estimate the underlying model parameters (Fig. 7)b. We generally found that all the model parameters were well estimated for the entire range of tested values. We found a small but systematic underestimation of the diffusion coefficient of the confined state that correlates with a slight overestimation of the confinement factor for datasets with lower lifetimes. The estimation of the shape parameter also worsened when the average lifetime is low.

### D. Inferring the number of motion states

In most biological SPT experiments, it is not fully clear how many states of motion should be expected (13, 40, 41). In such cases, a useful approach consists in evaluating the goodness of the model as a function of the number of parameters with Bayesian inference techniques (11). A simple yet efficient method to compute the number of states is the Bayesian Information Criterion (BIC). However, we demonstrated previously that the BIC tends to overestimate the number of states for large datasets due to inherent discrepancies between the model assumptions and the data (28). We found that adding a regularization term proportional to the log likelihood and the number of free parameters alleviates this issue.

To evaluate how well the number of states could be distinguished, we simulated a population of tracks with four distinct states of motion (immobile, directed, confined and diffusive) and transitions between states to test our model capability (see Fig. 9 to visualize a range of example tracks). Next, we designed an algorithm that starts by fitting the model with an overestimate of the number of states and then iteratively prunes the states that have the lowest impact on the maximum likelihood of the data. This results in a set of models whose likelihoods are a function of the number of states that increase until the correct number of states, and then plateaus for higher number of states, as exemplified in Fig. 8a (left). In this example, the BIC reaches a maximum at the true underlying number of states (Fig. 8a (left)). However, this is not universally true, as seen in Fig. S1, where the likelihood continues to increase when the number of states of the model passes the true underlying number of states. There, the regularization term recovers the true number of states. Additionally, we can see that the parameters estimated by the 4-state model are very close to the true underlying parameters (Fig. 8b-e).

**Fig. 8.**
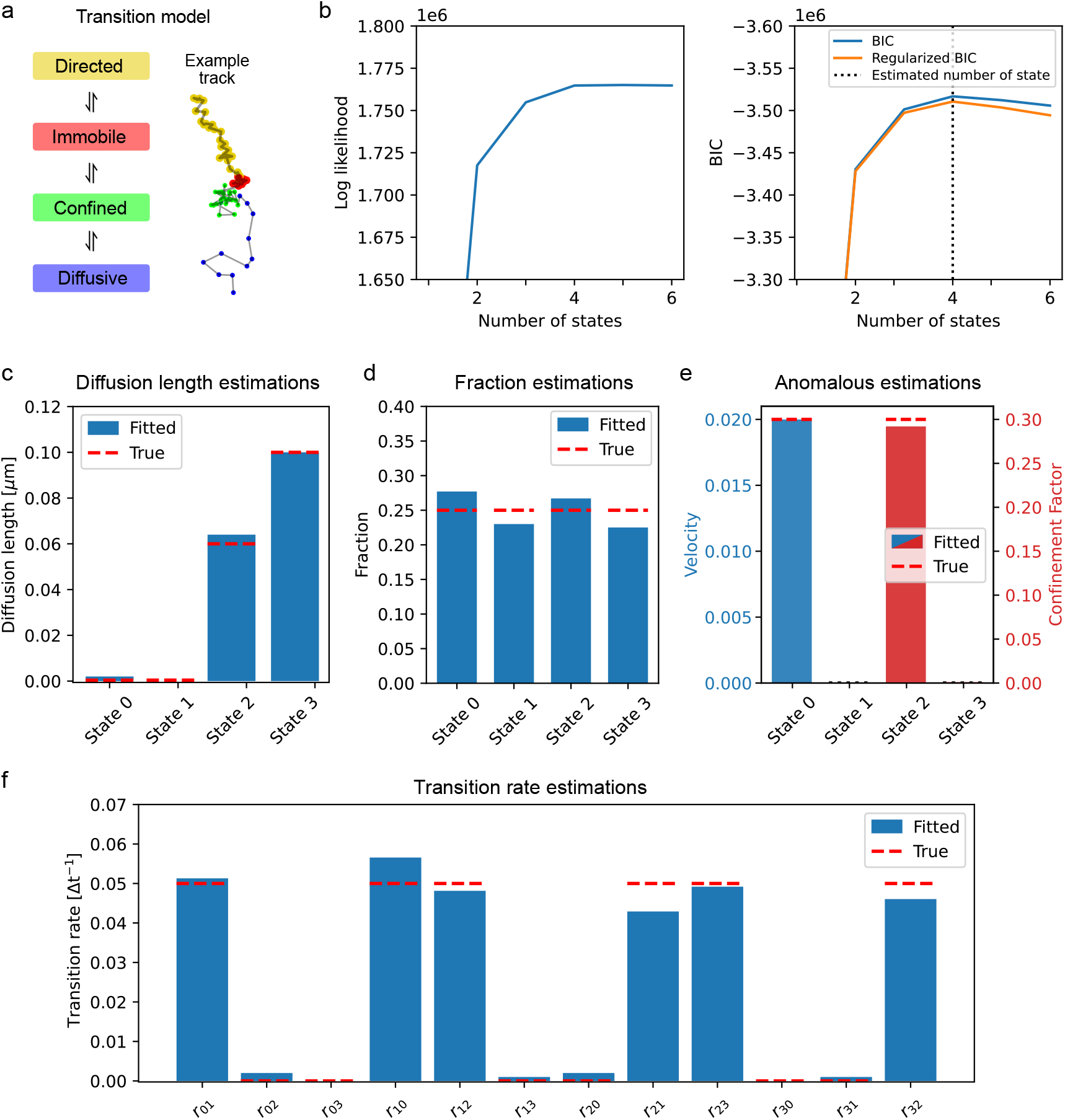
Estimating the number of states and model parameters from track populations. We simulated a population of 10,000 tracks of 60 time points in 4 states of motion (immobile, directed, confined and diffusive) to 1) test the capacity of our model to determine the number of states, 2) estimate the model parameters and 3) perform state predictions. **a**: Transition graph of the simulated tracks with an example track. The track is a made-up track made from segments of other tracks to ensure that the 4 states are present with little overlap between positions for maximal readability. **b**: Likelihood and Bayesian Information Criterion (BIC) of the model as a function of the number of states. The regularized BIC corresponds to the BIC supplemented with a regularization term equal to 0.0002 *k l* with *k* the number of parameters and *l* the likelihood of the model similarly to the term used in (24). **c-f**: Parameters estimated by the 4-state model. bars: estimated values, dashed lines: true underlying values.

**Fig. 9.**
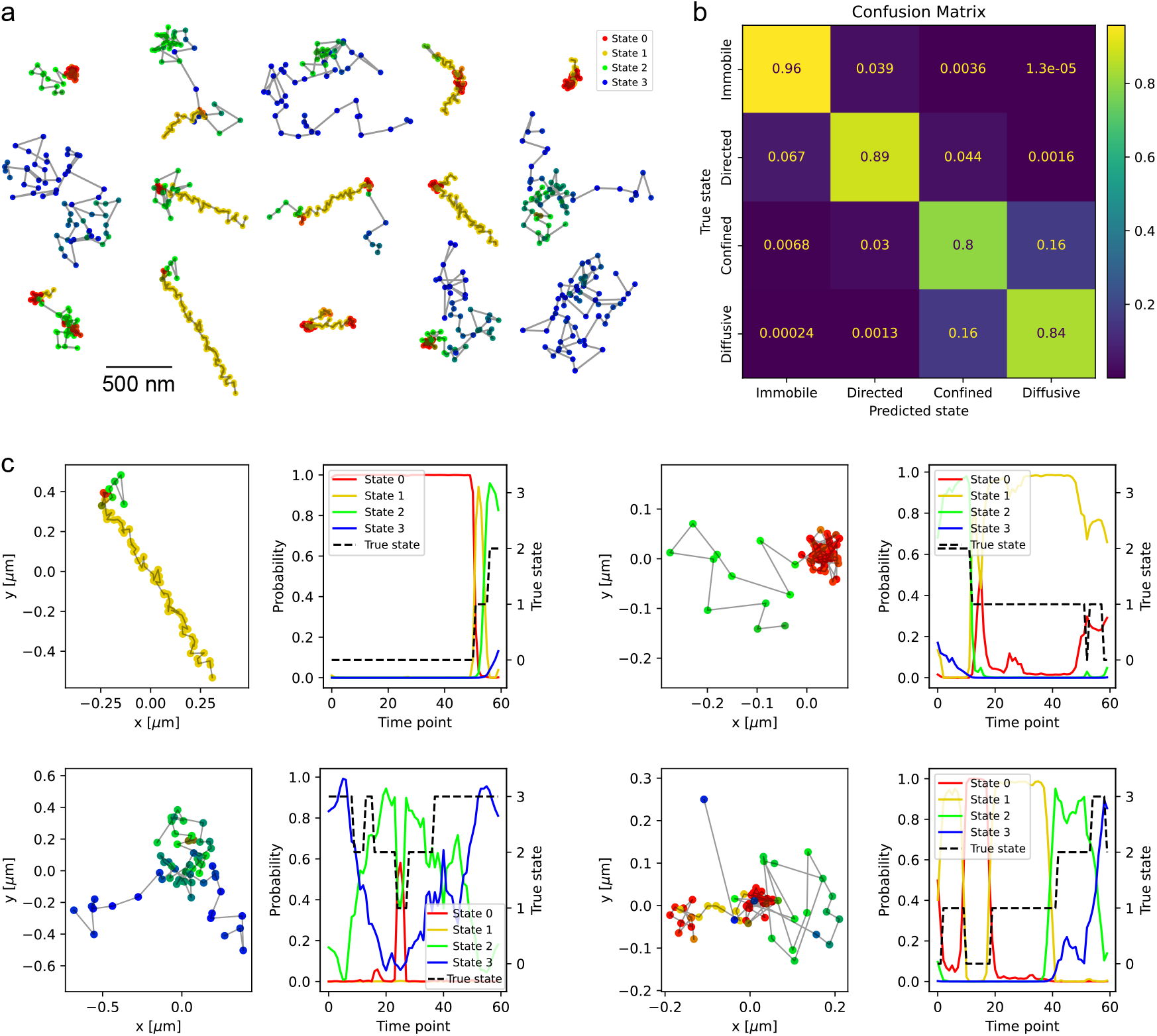
Intra-track state labeling. We used the tracks simulated for Fig. 8 and the 4-state model estimated from these tracks to assign a probability of being in the different states for each time point of the tracks. **a**: Random selection of tracks colored according to their state predictions. The pure colors (red for immobile, yellow for directed, green for confined and blue for diffusive) indicate a high probability of being in the corresponding state while mixed colors indicate intermediate probabilities. **b**: Confusion matrix of the state predictions of each time points of the 10,000 tracks for the 4 categories of motion. Categorical states for each time point were inferred based on the maximum state probability. **c**: 4 Examples of tracks from **a** (left sub-panels) represented next to the probabilities of being in each state through the tracks (right sub-panels).

### E. State predictions

Another important feature of ExaTrack is its capacity to classify which parts of each track are in the different states of motion. To do so, we compute the probability of each time point of a track to be in a given state of motion as was done for ExTrack (12). In Figure 9, we show the 4-state dataset and the model inferred in Fig. 8 that was used to compute the state probabilities along the tracks. Visual inspection of the tracks overlaid with their state probabilities (Fig. 9a,c) show the model is generally reliable at identifying the different states of motion and accurately depicts the transitions between motion states. The confusion matrix of the predicted states classified from the state probabilities (Fig. 9b) is a good quantitative indicator of how confident we can be on the annotated states. Instead of assigning a state to all the time points, we can also fix a threshold for state assignment to control the rate of false state labeling. Plotting the evolution of the state probabilities as a function of time next to a given track is a good way to obtain more insights into the performance of state labeling. Of course, for very transient states, annotations are much more difficult as shown in Fig. 9c.

## Discussion

Here, we presented ExaTrack, a tool designed to analyze particle tracks that contain multiple states of motion including immobilization, free diffusion, directed motion, confined diffusion with Markovian and gamma-distributed transitions between states. This tool has been tested over a wide range of simulated tracks to test the working range for reliably estimating the characteristics of the different states of motion such as the diffusion coefficient, the velocity or the degree of confinement, as well as the kinetics of transitions between states, that are the transition rates (or lifetime) and the shape parameter.

ExaTrack was designed for complex types of motion and transition kinetics. Handling non-Brownian tracks usually requires longer segments per state than previous Brownian motion models (10–12). For instance, estimating the confinement radius requires a particle to explore the confinement space which generally takes many time steps, as shown in the aTrack article (24). With this limitation in mind, we chose to consider a limited number of sequences of states which scales better with the number of states at a small cost of accuracy (see the methods section for more details). While not incompatible with the overall approach, ExaTrack does not consider intermediate states of diffusion at transitions, which leads to underestimations of the binding rates when the rate is higher than 0.1 Δ*t*^−1^, in agreement with previous results (12). Similarly, ExaTrack does not consider the defocalization bias (12, 42) which occurs when fast moving particles regularly enter and leave the depth of field. Note that tracks defocalize proportionally to the diffusion. As a result, short tracks are enriched in fast diffusion states while long tracks are enriched in low diffusion states. Discarding short or long tracks can then lead to survivor biases.

While our model is designed to identify a wide range of motion parameters, some parameter regimes are inherently ambiguous as they result in the same motion patterns. For example, free Brownian motion can be obtained using both our directed motion model and confined model, when the velocity or the confinement factor is near zero. Similarly, a particle can be considered immobile either by negligible diffusion and velocity (lengths much smaller than the localization error) or by strong confinement to an immobile potential well. Note that, the fitting model does not provide insight into whether states are significantly directed or confined compared to a pure Brownian null hypothesis and an additional analysis step like cross-validation must be carried to address this question. Individual states also become increasingly ambiguous when the transitions become more frequent. In this regime, it is likely necessary to record data at higher frame rates (13).

In ExaTrack, considering gamma-distributed lifetimes provides an additional degree of freedom to the model. While the kinetics of a sequence of multi-order reactions as in oligomerization, molecular motor progression, or polymerases (15, 32, 34, 37, 43) is complicated, estimating the shape parameter provides an idea of how time-dependent is the process. Determining the time-dependency of transition kinetics poses a challenge in terms of track length. Indeed, to precisely measure time-dependent lifetime kinetics, we need to track particles for the full lifetime of the state. This requires tracks that are typically longer than the state lifetimes. An alternative to using gamma-distributed lifetimes, that is not used here, is to explicitly add sequences of memoryless substates to the transition model with the same motion parameters. This approach would better fit more complex transition models but raises questions on how to connect sub-states. For example, transitions between all sub-states could be possible or they could follow a specific sequence. In either case, such models would result in better fits of the lifetimes but precisely identifying each of the transition rates would be complicated if not impossible.

ExaTrack assumes each state has a set of corresponding underlying parameter values. However, local heterogeneity and cell-to-cell variability within a dataset inevitably result in some distribution of true underlying parameter values including diffusion coefficients and transition rates. Further, it is possible that the diffusion coefficient of a particle may exhibit some time-dependent variation due to local changes of viscosity or crowding (4, 44). Analogously, the number of photons emitted by a fluorophore during a time step is inherently random inducing variations in the localization error (45). In the ExTrack article (12), we found that assuming fixed underlying localization errors and diffusion coefficients had little impact on the predicted parameters. For localization error, a way to alleviate this issue is to directly estimate the error from the particle brightness rather than the measured motion. It is significantly harder to determine the number of biologically relevant states when the true underlying model has a distribution of underlying parameters. In such cases, the maximum likelihood would show either no plateauing or delayed plateauing as a function of the number of states. Several model states might then be required to account for each of the biologically relevant states. Adapting ExaTrack to a Bayesian hierarchical model that considers the probability distribution of the model parameters would alleviate that issue (22, 46).

To our knowledge, ExaTrack is the first tool that uses a fully probabilistic model to consider transitions between a wide variety of states of motion such as immobile, diffusive, directed and confined motion. The most relevant available alternatives are either based on Hidden Markov Models (HMMs) or on change-points methods based on machine-learning. Most HMMs assume that the observed variables (emissions) only depend on the type of motion (10, 11, 47), preventing them from capturing the relationships between the observed localizations, the real particle positions and the anomalous variable. The current change-point-machine-learning methods are mainly focused on single-tracks as reported in (19) and therefore lack a pooled population model that considers multiple states and the kinetics of transitions between states. Moreover, at present they rely on mathematical models where states are characterized by mean squared displacements (MSDs) that increase with *t*^*α*^ with time *t* and anomalous exponent *α*. Such fractal implementations seems tenuous in many biological contexts as the forces that rule the motion typically vary with the scales. Of course, more physics-informed motion models, such as the ones used here, can be used for training.

In conclusion, ExaTrack is a versatile single-particle tracking analysis tool suited to analyze the dynamics of particles able to transition between a wide range of states of motion, namely, diffusion, directed and confined motion. This represents a novel set of particle tracking capabilities for tackling memory-dependent transitions between states while also accounting for localization error, diffusion, directed motion with changes of velocity or confined motion with changes of the confinement area. While we have thoroughly tested ExaTrack on a wide range of simulated data, its use remains to be demonstrated in real experiments. We believe that integrating probabilistic frameworks and the capacity to tackle both complex types of motion and transitions between states represents a major milestone for SPT analysis, improving the study of biological systems that often contain these features. Notably, we believe that this alternative to estimating the anomalous exponent offers a more biologically interpretable model for non-Brownian behavior.

## ACKNOWLEDGEMENTS

We wish to thank Claudiu Gradinaru, Xiaohan Zhou, Maria Shemeteva, Séamus Holden, Daniel Zenklusen, Om Mattagajasingh, Justina Chu and Sven van Teeffelen for helpful discussions. This project received funding from the European Union’s Horizon Europe research and innovation program under the Marie Skłodowska-Curie Postdoctoral Fellowship No. 101210381 (awarded to F.S.); the Natural Sciences and Engineering Research Council of Canada (NSERC Discovery grants RGPIN-2022-05142 to L.E.W.). L.E.W. is the recipient of a salary award from the Fonds de Recherche du Québec – Santé (FRQS) and acknowledges support from the Canada First Research Excellence Fund through the institut TransMedTech.

## Methods

### F. Gamma-distributed state transitions

To model gamma-distributed transitions, we use a rate matrix and a shape matrix that determine the transition rates and shape of the gamma distributions from state *s*_*i*_ to state *s*_*i*+1_ if 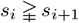. From these matrices, we infer the transition probability matrix given the current duration of the state. As the initial state duration is unknown, we assume the first transition to be independent of time. This method can therefore only work if multiple transitions can be observed within tracks.

### G. Moment matching approximation

In ExaTrack, the probability of a track given a sequences of states is a Gaussian of the track positions that we can compute with our recurrence methods. Therefore, the probability of a track is a mixture of Gaussians whose number of components increases with the number of states *n* to the power of the track length. To make this computation tractable even for long tracks, at each time step, we approximate the Gaussians that represent multiple sequences of states with similar properties into single Gaussians of averaged mean and variance. More specifically, we group the Gaussians of all the sequences of states that just transitioned into state *s*_*i*_ into one Gaussian. Then, we also group sequences of states that did not transition for *m* steps. This approach limits the number of elements of the Gaussian mixture to the number of states times *m*. Applying these approximations, for a 2-state model at a given time step *i* with *m* = 3, the memorized sequences […, s_i−3_, s_i−2_, s_i−1_, s_i_] are:

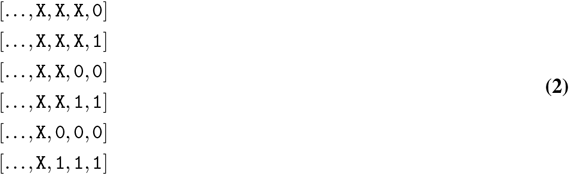

with *X* an undetermined state that cannot be the same as the following state if followed by a determined state. To enable gamma-distributed transitions, we keep track of the segment length and perform weighted averages of the segment lengths when merging sequences. At transitions, the current value of the anomalous variable that corresponds to the state at the previous step becomes irrelevant. We therefore integrate over this anomalous variable and initialize a new anomalous variable that corresponds to the new state of motion. This moment matching approximation comes at a cost of accuracy on the computed likelihood and model parameters. That cost is negligible in most of the tested datasets but can become relevant when multiples states have very similar motions and high transition rates. NB: during a given step we consider all the potential combinations of each of the sequences with an additional state so the maximum number of sequences of states to consider is (*n* + 1)^2^*m*.

### H. Modeling confinement

In this article, we assume that confinement results from the attraction of the particle of position *r* towards the center of a potential well of position *h* according to a confinement factor *l*. In a discrete process, this can be approximated by a first Brownian motion sub-step that updates the particle position: 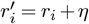 with *η* a normally distributed random variable of mean 0 and standard deviation 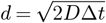and a second sub-step of attraction towards the potential well 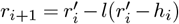. As a result, the discrete confinement factor that we estimate takes values between 0 and 1. However, real-life motion is continuous and this motion is described by the Ornstein–Uhlenbeck process (48) of stochastic differential equation *dx*_*t*_ = *θ*(*µ− x*_*t*_) + *σdW*_*t*_ with *W*_*t*_ the Wiener process. To compare our discrete model to the continuous reality, we then need to draw the relationship *l ≈* 1− *exp*(− *θt*) to compute the *θt* that must be used in our continuous-time simulations. Then, the confinement radius defined as the standard deviation of the distance between the real particle position and the center of the potential well can be computed as 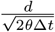. One of our conditions tests the influence of a confinement factor of 0.99, this translates into a continuous time confinement factor *θ*Δ*t* of around 4.605 and a confinement radius of *≈* 0.033 µm.

### I. Determining the type of motion

To determine the type of motion for a model with a given number of states, we first assign a type of motion to the different states. We then fit the model. Next, we swap the type of motion of one of the state and fit the model again. Thereafter, we compare the likelihoods of the two models and pick the model with the highest likelihood. We repeat this process for each of the states.

### J. Inferring the number of states

To determine the type of motion for a model with an unknown number of states, we start fitting a model with a large overestimate of the number of states. We then compute the likelihood of the model and estimate the impact of each of the states by removing them individually and computing the likelihood of the pruned model. Next, we select the pruned model with the highest likelihood, ensuring the pruning of the least relevant state and fit the pruned model. We repeat the pruning until the model possesses a unique state. The number of states can then be either manually determined by estimating the start of the likelihood plateau or by using the BIC or the Regularized BIC. If the estimated number of states in close to the initial number of states, restarting the procedure with a higher initial number of states might be required. Also, this algorithm does not swap the types of motion as it assumes overestimates of the initial number of states for both types of motion. Therefore, it is good practice to verify that the initial number of confined and directed states are both superior to the inferred numbers of confined and directed states. If more states are obtained than the number of biologically relevant states, the model states that capture the same biological state might be fused into a state of average parameters (and the model retrained) or kept as is while adding their fractions to quantify the biological state.

## Supplementary Figures

**Supplementary Figure 1.**
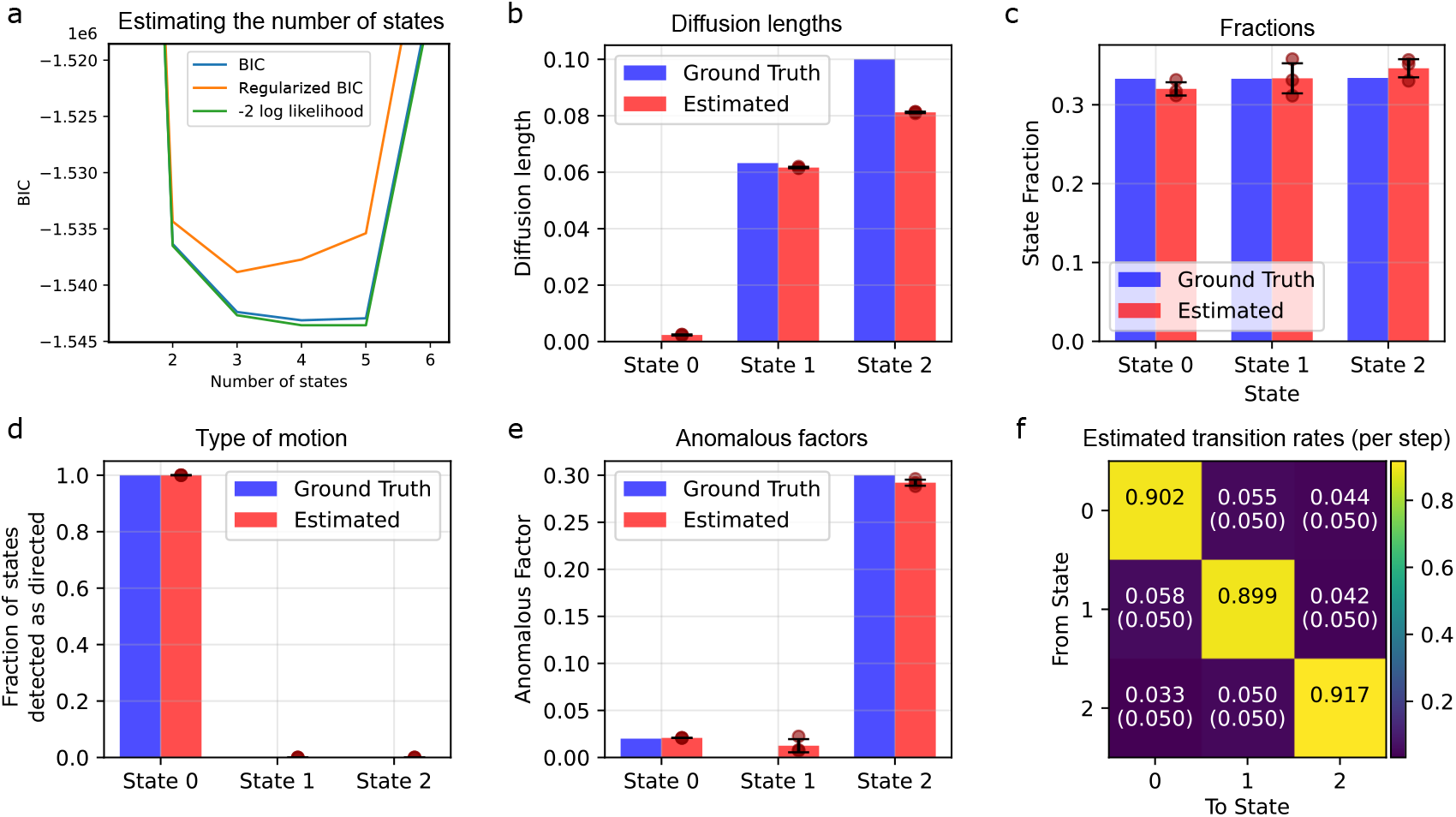
Estimating the number of states (3-state model). We simulated populations of 10,000 tracks of 30 time points in 3 states of motion (directed, confined and diffusive) to test the capacity of our model to determine the number of states and estimate the model parameters. **a** Likelihood, Bayesian Information Criterion (BIC) and regularized BIC (with the same penalization term 0.0002 *k l* as in Fig. 8). **b-e**: Estimations of the 3-state model parameters (diffusion length, fraction, type of motion and anomalous factors). In this version of the tool, the model automatically alternates the types of motion to determine the best suited motion type of each state. For the states 0, the anomalous factor then represents the velocity, while it represents the confinement factor for states 1 and 2. **f**: Matrix of the estimated transition rates from state *i* (rows) to state *j* (columns). The numbers in parentheses indicate the true transition rates.

## Bibliography

1. Megan A Steves, Changdong He, and Ke Xu. Single-molecule spectroscopy and super-resolution mapping of physicochemical parameters in living cells. Annual Review of Physical Chemistry, 75, 2024.

2. François Laurent, Charlotte Floderer, Cyril Favard, Delphine Muriaux, Jean-Baptiste Masson, and Christian L Vestergaard. Mapping spatio-temporal dynamics of single biomolecules in living cells. Physical biology, 17(1):015003, 2019.

3. Gavin Schlissel, Miram Meziane, Domenic Narducci, Anders S Hansen, and Pulin Li. Diffusion barriers imposed by tissue topology shape hedgehog morphogen gradients. Proceedings of the National Academy of Sciences, 121(36):e2400677121, 2024.

4. José Losa, François Simon, Dmitrii Linnik, Saniye Gül Kaya, Marc CA Stuart, Artem Stetsenko, Rinse de Boer, Franz Y Ho, Danny Incarnato, Jan Stevens, et al. Condition-dependent, amorphous protein agglomerates control cytoplasmic rheology. bioRxiv, pages 2025–06, 2025.

5. Limin Xiang, Kun Chen, Rui Yan, Wan Li, and Ke Xu. Single-molecule displacement mapping unveils nanoscale heterogeneities in intracellular diffusivity. Nature methods, 17(5):524–530, 2020.

6. Antoine Vigouroux, Baptiste Cordier, Andrey Aristov, Laura Alvarez, Gizem Özbaykal, Thibault Chaze, Enno Rainer Oldewurtel, Mariette Matondo, Felipe Cava, David Bikard, et al. Class-a penicillin binding proteins do not contribute to cell shape but repair cell-wall defects. Elife, 9:e51998, 2020.

7. Janka Zsok, Francois Simon, Göksu Bayrak, Luljeta Isaki, Nina Kerff, Yoana Kicheva, Amy Wolstenholme, Lucien E Weiss, and Elisa Dultz. Nuclear basket proteins regulate the distribution and mobility of nuclear pore complexes in budding yeast. Molecular Biology of the Cell, 35(11):ar143, 2024.

8. Chieh-Teng Cheng, Jye-Chian Hsiao, Alexander Hoffmann, and Hsiung-Lin Tu. Tnfr1 mediates heterogeneity in single-cell nf-κb activation. Iscience, 27(4), 2024.

9. Tomáš Janovič, Gloria I Perez, and Jens C Schmidt. Trf1 and trf2 form distinct shelterin subcomplexes at telomeres. bioRxiv, 2024.

10. Raibatak Das, Christopher W Cairo, and Daniel Coombs. A hidden markov model for single particle tracks quantifies dynamic interactions between lfa-1 and the actin cytoskeleton. PLoS computational biology, 5(11):e1000556, 2009.

11. Fredrik Persson, Martin Lindén, Cecilia Unoson, and Johan Elf. Extracting intracellular diffusive states and transition rates from single-molecule tracking data. Nature methods, 10(3):265, 2013.

12. François Simon, Jean-Yves Tinevez, and Sven van Teeffelen. Extrack characterizes transition kinetics and diffusion in noisy single-particle tracks. Journal of Cell Biology, 222(5):e202208059, 2023.

13. François Simon, Lucien E Weiss, and Sven van Teeffelen. A guide to single-particle tracking. Nature Reviews Methods Primers, 4(1):66, 2024.

14. Amy N Moores and Stephan Uphoff. Robust quantification of live-cell single-molecule tracking data for fluorophores with different photophysical properties. The Journal of Physical Chemistry B, 128(30):7291–7303, 2024.

15. Chunte Sam Peng, Yunxiang Zhang, Qian Liu, G Edward Marti, Yu-Wen Alvin Huang, Thomas C Suedhof, Bianxiao Cui, and Steven Chu. Nanometer-resolution tracking of single cargo reveals dynein motor mechanisms. Nature Chemical Biology, 21(5):648–656, 2025.

16. Paolo Pierobon, Sarra Achouri, Sébastien Courty, Alexander R Dunn, James A Spudich, Maxime Dahan, and Giovanni Cappello. Velocity, processivity, and individual steps of single myosin v molecules in live cells. Biophysical journal, 96(10):4268–4275, 2009.

17. Ralf Metzler, Jae-Hyung Jeon, Andrey G Cherstvy, and Eli Barkai. Anomalous diffusion models and their properties: non-stationarity, non-ergodicity, and ageing at the centenary of single particle tracking. Physical Chemistry Chemical Physics, 16(44):24128–24164, 2014.

18. Felix Höfling and Thomas Franosch. Anomalous transport in the crowded world of biological cells. Reports on Progress in Physics, 76(4):046602, 2013.

19. Gorka Muñoz-Gil, Harshith Bachimanchi, Jesús Pineda, Benjamin Midtvedt, Gabriel Fernández-Fernández, Borja Requena, Yusef Ahsini, Solomon Asghar, Jaeyong Bae, Francisco J Barrantes, et al. Quantitative evaluation of methods to analyze motion changes in single-particle experiments. Nature Communications, 16(1):6749, 2025.

20. Jacob Kæstel-Hansen, Marilina de Sautu, Anand Saminathan, Gustavo Scanavachi, Ricardo F Bango Da Cunha Correia, Annette Juma Nielsen, Sara Vogt Bleshøy, Konstantinos Tsolakidis, Wouter Boomsma, Tomas Kirchhausen, et al. Deep learning-assisted analysis of single-particle tracking for automated correlation between diffusion and function. Nature Methods, pages 1–10, 2025.

21. Wenjie Cai, Yi Hu, Xiang Qu, Hui Zhao, Gongyi Wang, Jing Li, and Zihan Huang. Machine learning analysis of anomalous diffusion. The European Physical Journal Plus, 140(3):183, 2025.

22. Ziyuan Chen, Laurent Geffroy, and Julie Suzanne Biteen. Nobias: Analyzing anomalous diffusion in single-molecule tracks with nonparametric bayesian inference. Frontiers in bioinformatics, page 40, 2021.

23. Benjamin Recht, Rebecca Roelofs, Ludwig Schmidt, and Vaishaal Shankar. Do imagenet classifiers generalize to imagenet? In International conference on machine learning, pages 5389–5400. PMLR, 2019.

24. François Simon, Guillaume Ramadier, Inès Fonquernie, Janka Zsok, Sergiy Patskovsky, Michel Meunier, Caroline Boudoux, Elisa Dultz, and Lucien E Weiss. Detecting directed motion and confinement in single-particle trajectories using hidden variables. bioRxiv, pages 2024–04, 2024.

25. Jason Bernstein and John Fricks. Analysis of single particle diffusion with transient binding using particle filtering. Journal of theoretical biology, 401:109–121, 2016.

26. Paddy J Slator and Nigel J Burroughs. A hidden markov model for detecting confinement in single-particle tracking trajectories. Biophysical journal, 115(9):1741–1754, 2018.

27. Nilah Monnier, Zachary Barry, Hye Yoon Park, Kuan-Chung Su, Zachary Katz, Brian P English, Arkajit Dey, Keyao Pan, Iain M Cheeseman, Robert H Singer, et al. Inferring transient particle transport dynamics in live cells. Nature Methods, 12(9):838, 2015.

28. Francois C Simon and Justin Cardona. Fast analytical method to integrate multivariate gaussians over hidden variables. 2024.

29. Jochem NA Vink, Stan JJ Brouns, and Johannes Hohlbein. Extracting transition rates in particle tracking using analytical diffusion distribution analysis. Biophysical Journal, 119(10):1970–1983, 2020.

30. Charles R Sanders. Biomolecular ligand-receptor binding studies: theory, practice, and analysis. Nashville: Vanderbilt University, pages 1–43, 2010.

31. Wylie Stroberg and Santiago Schnell. On the validity and errors of the pseudo-first-order kinetics in ligand–receptor binding. Mathematical Biosciences, 287:3–11, 2017.

32. Athel Cornish-Bowden. Fundamentals of enzyme kinetics. John Wiley & Sons, 2013.

33. Daniel L Floyd, Stephen C Harrison, and Antoine M Van Oijen. Analysis of kinetic intermediates in single-particle dwell-time distributions. Biophysical journal, 99(2):360–366, 2010.

34. Sridharan Rajagopalan, Fang Huang, and Alan R Fersht. Single-molecule characterization of oligomerization kinetics and equilibria of the tumor suppressor p53. Nucleic acids research, 39(6): 2294–2303, 2011.

35. David M Kanno and Marcia Levitus. Protein oligomerization equilibria and kinetics investigated by fluorescence correlation spectroscopy: a mathematical treatment. The Journal of Physical Chemistry B, 118(43):12404–12415, 2014.

36. Samara L Reck-Peterson, Ahmet Yildiz, Andrew P Carter, Arne Gennerich, Nan Zhang, and Ronald D Vale. Single-molecule analysis of dynein processivity and stepping behavior. Cell, 126(2): 335–348, 2006.

37. Kristina M Herbert, Arthur La Porta, Becky J Wong, Rachel A Mooney, Keir C Neuman, Robert Landick, and Steven M Block. Sequence-resolved detection of pausing by single rna polymerase molecules. Cell, 125(6):1083–1094, 2006.

38. Aurelien Pelissier, Miroslav Phan, Niko Beerenwinkel, and Maria Rodriguez Martinez. Practical and scalable simulations of non-markovian stochastic processes. arXiv preprint arXiv:2212.05059, 2022.

39. M Weiss. Use of gamma distributed residence times in pharmacokinetics. European journal of clinical pharmacology, 25(5):695–702, 1983.

40. Alec Heckert, Liza Dahal, Robert Tjian, and Xavier Darzacq. Recovering mixtures of fast-diffusing states from short single-particle trajectories. Elife, 11:e70169, 2022.

41. Chiara Schirripa Spagnolo and Stefano Luin. Trajectory analysis in single-particle tracking: From mean squared displacement to machine learning approaches. International Journal of Molecular Sciences, 25(16):8660, 2024.

42. Anders S Hansen, Maxime Woringer, Jonathan B Grimm, Luke D Lavis, Robert Tjian, and Xavier Darzacq. Robust model-based analysis of single-particle tracking experiments with spot-on. Elife, 7:e33125, 2018.

43. Keith J Mickolajczyk and William O Hancock. Kinesin processivity is determined by a kinetic race from a vulnerable one-head-bound state. Biophysical journal, 112(12):2615–2623, 2017.

44. Wan Li and Ke Xu. Super-resolution mapping and quantification of molecular diffusion via single-molecule displacement/diffusivity mapping (sm d m). Accounts of Chemical Research, 58(8): 1224–1235, 2025.

45. Yan Chen, Joachim D Müller, Peter TC So, and Enrico Gratton. The photon counting histogram in fluorescence fluctuation spectroscopy. Biophysical journal, 77(1):553–567, 1999.

46. Jan-Willem Meent, Jonathan Bronson, Frank Wood, Ruben Gonzalez Jr, and Chris Wiggins. Hierarchically-coupled hidden markov models for learning kinetic rates from single-molecule data. In International Conference on Machine Learning, pages 361–369. PMLR, 2013.

47. Martin Lindén, Vladimir Ćurić, Elias Amselem, and Johan Elf. Pointwise error estimates in localization microscopy. Nature communications, 8(1):1–9, 2017.

48. Leonard Salomon Ornstein. On the theory of the brownian motion. Physical review, 36:823–841, 1930.

